# Improved spatial ecological sampling using open data and standardization: an example from malaria mosquito surveillance

**DOI:** 10.1101/465963

**Authors:** Luigi Sedda, Eric R. Lucas, Luc S. Djogbénou, Ako V.C. Edi, Alexander Egyir-Yawson, Bilali I. Kabula, Janet Midega, Eric Ochomo, David Weetman, Martin J. Donnelly

**Affiliations:** Centre for Health Information, Computation and Statistics (CHICAS), Lancaster Medical School, Furness Building, Lancaster University, Lancaster, LA1 4YG, UK; Department of Vector Biology, Liverpool School of Tropical Medicine, Pembroke Place, Liverpool L3 5QA, UK; Institut Régional de Santé Publique/Université d’Abomey–Calavi, Ouidah, Benin; Centre Suisse de Recherches Scientifiques en Cote d’Ivoire, 01 BP 1303 Abidjan 01, Cote d’Ivoire; Department of Biomedical Sciences, University of Cape Coast, Cape Coast, Ghana; National Institute for Medical Research (NIMR), Amani Centre, P.O. Box 81, Muheza, Tanzania; Centre for Geographic Medicine Research, Kenya Medical Research Institute, P.O. Box 230, 80108 Kilifi, Kenya.; Centre for Global Health Research, Kenya Medical Research Institute, Kisumu, Kenya.

**Author notes:** Wellcome Sanger Institute, Hinxton, Cambridge CB10 1SA, UK.

**Keywords:** Mosquito sampling, stratification, lattice sampling design, model-based geostatistics, Sub-Saharan Africa

## Abstract

Vector-borne disease control relies on efficient vector surveillance, mostly carried out using traps whose number and locations are often determined by expert opinion rather than a rigorous quantitative sampling design. In this work we first propose a framework for ecological sampling design which in its preliminary stages can take into account environmental conditions obtained from open data (i.e. remote sensing and meteorological stations). These environmental data are used to delimit the area into ecologically homogenous strata. By employing a model-based sampling design, the traps are deployed among the strata using a mixture of random and grid locations which allows balancing predictions and fitting accuracies. Sample sizes and the effect of ecological strata on sample sizes are estimated from previous sampling campaigns. Notably, we found that a configuration of 30 locations with 4 households each (120 traps) will have a similar accuracy in the estimates of mosquito abundance as 300 random samples. In addition, we show that random sampling independently from ecological strata, produces biased estimates of the mosquito abundance. Finally, we propose standardizing reporting of sampling designs to allow transparency and repetition / re-use in subsequent sampling campaigns.

## Introduction

Sampling design is a crucial step in any survey as it affects the quality of data collection and analysis (*1*). Sampling strategies should therefore be designed to maximise the effectiveness of the study, using any relevant preliminary and background data available (*2*). Furthermore, because published sampling strategies frequently inspire designs for future studies, both the design details and justification should be rigorously reported. Despite improvement in recent years, both the use of available informative data and the rigour with which sampling designs are reported continue to fall short of what could be achieved (*3*). Specifically, the amount of environmental data available from open-data platforms is often acknowledged but rarely exploited to support sampling design, while the necessary information for study repeatability, comparability or usability are often inadequately reported. Such data can support representativeness in investigations of population dynamics, epidemiological processes, and biological studies. Here we use an example from malaria vector surveillance to design a sampling strategy for collecting mosquitoes for whole genome sequencing based monitoring and evaluation. Genomic technologies are radically transforming our understanding of vector-borne disease transmission dynamics (*4*) due to the capacity to unveil complex interaction between human, pathogen, vector and environment. Whole genome sequencing projects have revealed novel genetic loci associated with increased susceptibility to malaria in the human host (*5, 6*) and made major contributions to our understanding of how anti-malarial and insecticide resistance evolves (*6*). However, the impact of environment on genotype distributions is much more poorly understood, reflecting at least in part the use of insufficiently ecologically-informed sampling strategies. Much of the sampling conducted in vector surveillance studies is opportunistic and lacks a rigorous sampling framework. Often, ecological and entomological sampling designs rely solely on resource availability rather than aiming to maximize representativeness and precision of the variable of interest, e.g. collectors target locations where disease vectors are known to be abundant.

Designing a field sampling strategy requires three decisions: what is the variable of interest (formally the estimator, e.g. vector density), the sampling approach (e.g. model-based or not, in other words, whether we apply a model to choose the sampling location or just use pre-existing knowledge) and sampling location distribution (e.g. the number and spatial / temporal distribution of sampling points). These decisions constitute the sampling strategy trinity (*7*) in which each element strictly depends on the other two. Sampling strategies are further complicated by deterministic (e.g. due to age, environment, socio-economic, etc) and stochastic (spatio-temporal autocorrelation) factors. Our literature search in Web of Science on spatial sampling of mosquitoes (search terms: mosquito or anopheles AND sampling AND spatial, in title/keywords/abstract) shows that while all studies provide a general description of the sampling design, only a limited number of papers (i.e. (*8-11*)) give a detailed description of the rationale, decisions and calculations related to the “Where, When, How and How many” samples to collect (see for example the reviews from (*12*) and (*13*)). In other literature, partial justification of the sampling design is provided. For example, (*14-22*) used previous surveillance information and remote sensing data to identify potential mosquito habitat types (or, in statistical terms, “strata”). However, the method used or assumptions made to choose the within-strata location and number of traps were not described, perhaps because these were entirely guided by practical considerations (i.e. (*23*)). Conversely, descriptions of sampling over time are often provided in detail, with explicit information on the frequency and length of the sampling campaign.

The picture that emerges from the literature is that using habitat stratification to inform sampling is a common procedure in vector biology, but often based on subjective or qualitative decisions. However, stratification has a fundamental role in describing and reducing the error in estimates of mosquito variation, which in turn influences surveillance success, assessment of epidemiological risk and genetic diversity (*24*). Quantitative stratification is usually performed by identifying a set of (independent) environmental variables that can be used to define strata within which the property or properties under study (i.e. insecticide resistance) is/are relatively homogeneous (*13*). Unless the spatial or spatiotemporal process (i.e. the spatial or spatiotemporal autocorrelation of the property under study) is tested and found negligible (*25*), these approaches often incorrectly assume independence between samples in space and time (*26*) (an unrealistic assumption for most of the ecological processes). Spatial and spatiotemporal heterogeneity can be accounted for in sampling design by adopting a geostatistical model-based sampling design (*8, 27*).

Ecological stratification of sampling designs is now facilitated by web-based open data providers, allowing rapid access to large amounts of information on climate and land-use, which are commonly associated with biogeographic patterns of human and animal health and species distribution (*28*). This availability of open data (largely remote sensing) for almost every global location, combined with appropriate spatiotemporal algorithms (*15*), make quantitative ecological stratification more accessible as a preliminary step to any sampling programme. Nevertheless “*very few studies propose, at an early phase of research work, objective sampling strategies that are consistent with both study goals and constraints*” (*13*).

In this work we propose a framework for optimising the sampling design of the spatial distribution of mosquito populations using open data, which we hope will be relevant to a wide range of ecological, disease monitoring and genomic studies. The open data are used to ecologically-characterise the area(s) under study and inform the location of each trap or in general collection point. The sample size is calculated based on previous mosquito surveys and sample locations are defined to balance prediction and parameterization, i.e. the accuracy in predictions and the goodness of model fitting. The effect of ecological strata on sampling size is estimated from a previous malaria control surveillance campaign. Finally, we discuss the necessity and benefits of a standardization of the sampling design procedures and reports to make them repeatable and reusable.

## Materials

### GAARDian project

The sampling design described in this work has been developed within the UK-MRC-funded GAARDian project (https://www.anophelesgenomics.org/gaardian). The main objective of this project is to investigate the spatial and temporal scale of variation in mosquito genomes to improve our understanding of the processes underlying the spread of insecticide resistance. Insecticide resistance is a major threat to the sustained control of malaria, as 260 million averted clinical cases of malaria have been due to the use of insecticides that target the mosquito vector (*29*).

### Study area

Six sites were chosen based on suspected use of insecticide or presence of insecticide resistance (Fig. 1) (see below description for each site). Around each site, an operable area was determined as the largest area where traps can be deployed and routinely checked by two operators, within a 60 × 60km square centred on the site.

**Figure 1.**
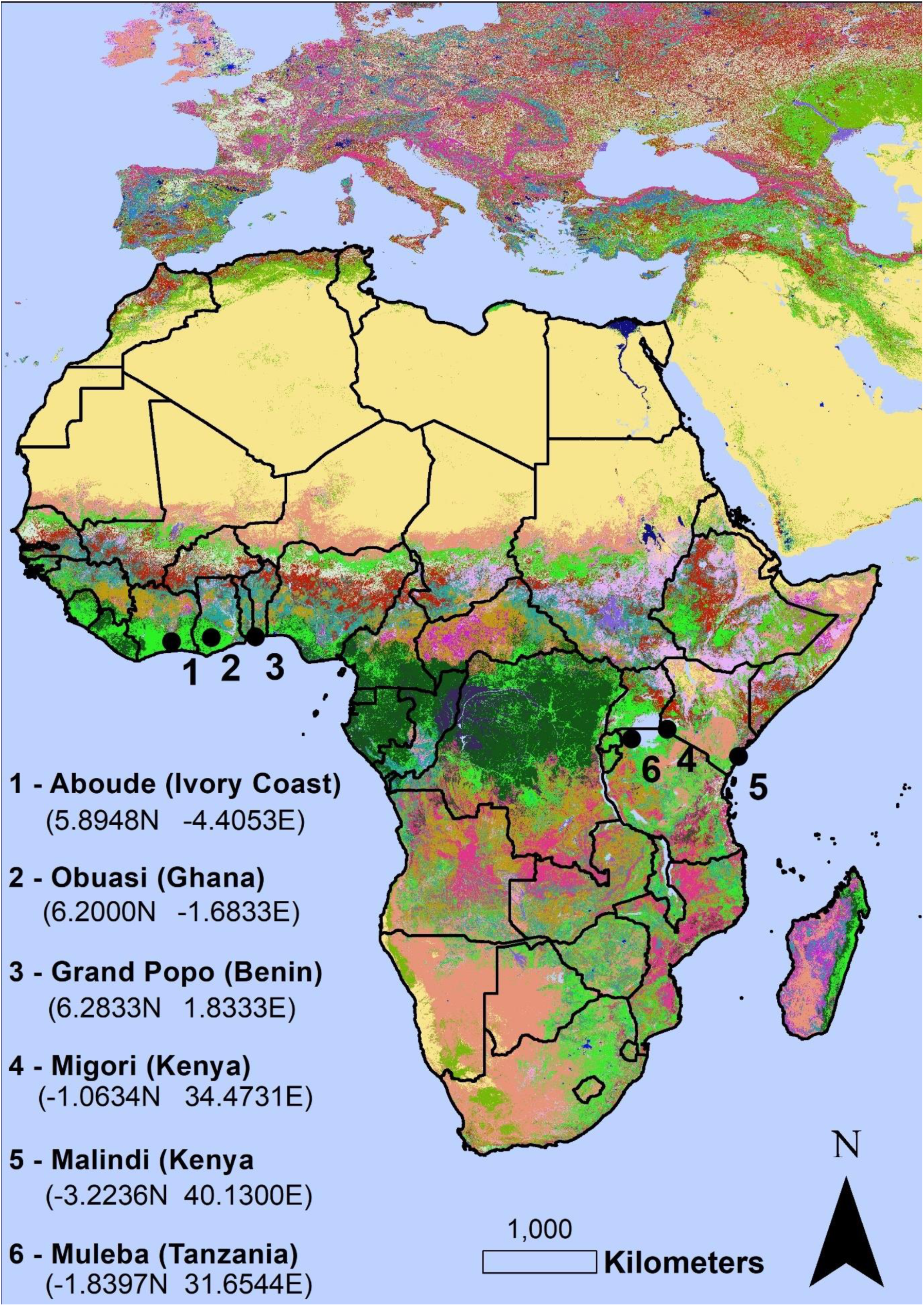
Location of the GAARDian sampling sites, shown on a land cover background (GlobeLand30 land covers). Map was made using ArcMap 10.4 (http://desktop.arcgis.com/en/arcmap/). Source administrative limits: http://www.maplibrary.org/library/index.htm.

#### Migori

Migori, is located in western Kenya, about 50km from Lake Victoria and with elevation ranging from 1200m to 1500m above sea level. The average annual temperature is 21°C, average relative humidity is 65% with average annual rainfall of 1,000 -1,800mm. The area experiences long rains from April to June and short rains from September to October. The land is mainly used for cultivation and grazing. There are some studies on malaria burden from the area (*30, 31*), but none on mosquito abundance or insecticide resistance even if indoor residual spraying is taking place (see methods).

#### Aboude

This area is located between Aboude and Agboville villages, in Southern Côte d’Ivoire. Aboude is located in the evergreen forest zone with altitude between 30 and 100 m above sea level. The climate is divided into four seasons: a long rainy season (April-July), a short dry season (August-September), a short rainy season (October-November) and a long dry season (December to March). Average temperature is around 27°C and average rainfall of 120mm. Relative humidity ranges from 70 to 85%. The hydrographic network of the region is very diversified and characterized by the presence of the Bandama and the N′zi Rivers with several streams. The primary activity of the rural population is agriculture with mainly cocoa, rubber, vegetable and irrigated rice fields with large use of pesticides. Malaria transmission occurs during the rainy seasons, between April and November (*32*) but insecticide resistance has not been documented to date.

#### Grand Popo

The study site in Benin is in the southwestern coastal part of the country. Elevation ranges from 0m to 70m above sea level. The average temperature is 28.9°C, average relative humidity is 76% with average annual rainfall of 190mm. The rainy season is characterized by abundant rains during April to July, and a lower amount of rain from September to October. The area is mostly urban and cultivated, and use of pesticide is common. Studies have been published on malaria incidence and bednet use (*33-35*), however there are no studies on mosquito species distribution or insecticide resistance.

#### Obuasi

Obuasi is located in the southern part of the Ashanti region of Ghana about 64 km south-west of the regional capital Kumasi. The area has an undulating terrain with most of the hills rising above 500 meters above sea level and vegetation characteristic of the moist semi-deciduous forest type. The climate is semi-equatorial and characterised by two rainy seasons. The first season starts from March and ends in July and the second from September to November. The mean annual rainfall ranges between 125mm and 175 mm, while the mean average annual temperature is 25.5 °C and relative humidity 75% – 80% in the wet season. Agricultural activities in the area include crop farming, livestock rearing, tree planting and fish farming. Mining and quarry forms the second largest industrial activity in the municipality and creates potential mosquito breeding sites all year round. Resistance to multiple insecticides in Obuasi has been documented in *Anopheles gambiae and An. funestus* mosquitoes (*36-38*).

#### Malindi

The study site contains the large town of Malindi with approximately 210,000 inhabitants. The climate is tropical, with a cooler season from June to September, with daytime temperatures around 27-28 °C, and a hotter and humid season from November to April, with daytime temperatures above 30 °C. Relative humidity ranges between 80-85%. Malindi is comprised of commercial and residential areas, agricultural and undeveloped areas, and hotels and stores along the coast. Tourism, retail, fishing, and trading are the major economic activities. This area is within Kenya’s endemic malaria zone with all-year risk of malaria transmission (*39*). The major malaria control intervention in Malindi is the use of pyrethroid treated bednets. Studies to detect insecticide resistance show suspected *Anopheles* resistance to pyrethroids (*40, 41*).

#### Muleba

Muleba is in the Kagera region of northwest Tanzania on the western shore of Lake Victoria. The district lies at 1100-1600m above sea level. There are two rainy seasons: “long rains” in March – June (average monthly rainfall 300 mm) and “short rains” in October-December (average monthly rainfall 160 mm). Average annual temperature is 21°C (with minimum- maximum range of 15°C-28°C) and average relative humidity of 66%. The area is mainly rural and is used for agriculture. Malaria transmission occurs throughout the year and peaks after the rainy seasons. The predominant malaria vectors are *Anopheles gambiae s.s*. and *An. arabiensis*, in which pyrethroid resistance has been detected (*42*).

### Environmental data

We used open data information from several sources to stratify the ecological variations of each study site. These data include land cover, climate and topography.

GlobeLand30 (http://www.globallandcover.com/GLC30Download/index.aspx) is a global land cover map of 30m resolution produced by the National Geomatics Center of China and containing 10 land cover classes (full description of classes in (*43*)). The images used for GlobeLand30 classification are multispectral images, including the TM5 and ETM + of America Land Resources Satellite (Landsat) and the multispectral images of China Environmental Disaster Alleviation Satellite (HJ-1). GlobeLand30 raster adopts WGS84 coordinate system, UTM projection, 6-degree zoning and the reference ellipsoid is WGS 84 ellipsoid.

The moderate-resolution imaging spectroradiometer (MODIS) satellite products are provided in monthly time-series at 0.05 degree (~5km) resolution from observations by the MODIS sensor on Terra (AM) for the period February 2000 to December 2013 inclusive and available at (https://ora.ox.ac.uk/objects/uuid:896bf37f-a56b-4bc0-9595-8c9201161973) (*44*). The following MODIS products were used:

- MODIS Enhanced Vegetation Index (EVI) from the MOD13C2 product comprises monthly, global EVI. This resource provides consistent spatial and temporal comparisons of vegetation canopy greenness, a composite property of leaf area, quantity of chlorophyll and canopy structure. EVI improves sensitivity over dense vegetation conditions or heterogeneous landscapes when compared to Normalized Difference Vegetation Index (NDVI).
- MODIS Air Temperature (Temp) from the MOD07_L2 Atmospheric Profile product comprises monthly, global temperature at the closest level to the earth’s surface.
- MODIS Evapotranspiration (ET) from the MOD16 Global Evapotranspiration product is calculated monthly as the ratio of Actual to Potential Evapotranspiration (AET/PET).

Precipitation was obtained from WorldClim Version 2 as average annual precipitation from 1970 to 2000 at 30 arcseconds (1 km^2^ ca.) (http://worldclim.org/version2) (*45*). Finally, elevation was obtained from the NASA Shuttle Radar Topographic Mission (SRTM) 90m Digital Elevation Database v4.1. The SRTM 90m DEM’s have a resolution of 90m at the equator. The DEM is available in geographic coordinate system – WGS84 datum (https://drive.google.com/drive/folders/0B_J08t5spvd8VWJPbTB3anNHamc).

Land cover, precipitation and elevation were re-projected at 5km, the same spatial resolution of the MODIS products, using a nearest-neighbour method for the categorical variable (land cover) and the bilinear interpolation method for continuous variables (precipitation and elevation) (*46*).

## Methods

We have adapted the sampling framework proposed by Wang et al. (*7*) to include the identification of ecological strata:

i. Sample size optimization.
ii. Stratification (ecological delineation).
iii. Spatial allocation of the sampling households. For the present study the malaria vector species we are targeting, within the *An. gambiae* species complex, are usually highly anthropophilic and commonly found in houses.

Description of each step is given below.

### Sampling size optimization

For one of the collection areas (Migori, Kenya, location 4 in Fig. 1), additional data were available from entomological surveillance carried out from December 2015 to September 2017 as part of indoor residual spraying (IRS) (Abong’o et al unpublished; http://www.africairs.net/about-airs/), which we will refer to as AIRS data hereafter. Following the methodology of the GAARDian project, collections are made using attractant light traps, which by placement near sleeping space, sample female mosquitoes that are actively seeking a blood meal. Light trap collections can be an accurate proxy of transmission risk (*47*). We used this preliminary information about mosquito abundance to estimate the optimal sample size (in terms of mosquito distribution) to be used in all sites.

From the AIRS data, we first estimated the spatial covariance function (via maximum likelihood estimation, (*48*)) that was used to simulate a Log Gaussian Cox process (LGCP) (*49*) mimicking the mosquito distribution process found in Migori. This can be translated in lay words as a process (mosquito catches) that is environmentally driven but producing values of catches that can be considered independent (i.e. catch on one occasion does not predict subsequent catches in the same or nearby locations).

The Gaussian random field is of the form (*50*):

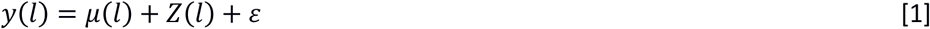

where *l* is the location, *μ* is the mean, **Z** is the gaussian process with Matern correlation function, and *ε* is the error term (noise or nugget).

The Matern correlation function has the general form:

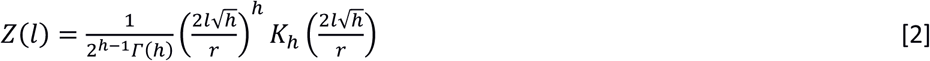

where **K**_*h*_(·) is the modified Bessel function of order *h* and *r* is the spatial range (*51*). Both *h* and *r* must be positive and different from 0.

Finally the Poisson LGCP can be written as (*52*):

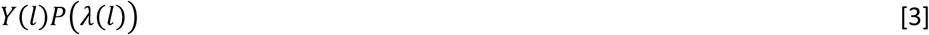

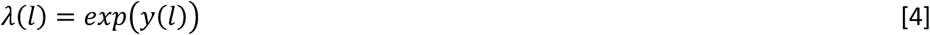

where *Y* is the mosquito density point process and *λ* is the conditional mean.

From the LGCP we predicted the estimated variance in the parameters of the spatial covariance function and the prediction error for a set of sample sizes (15, 30, 75, 150, 200 and 300) assumed randomly allocated in the area of Migori.

This will allow the allocation of the (limited) resources to obtain the sample size that will produce the desired prediction error (which should be lower than the expected average number of mosquitoes caught) and variance in the spatial covariance parameters (if this is an objective of the sampling design).

### Stratification (ecological delineation)

In many areas of physical, engineering, life and social sciences, inferential and predictive classification are prevalent tools to discriminate between classes and to interpret the differences. Examples range from identification of ecological niches to brain and bone anomalies. While the growing amount of open access information enables discrimination among a large number of ecological classes, many traditional algorithms fail for these data because of decreased classification performance (leading to overfitting) and mathematical/practical limitations (*53*). One method is to describe ecological strata in terms of transformed environmental variation (i.e. factorial analyses) (*13*), but the results can be difficult to interpret. By contrast, discriminant analysis (DA) requires less computational time and resources because no parameter tuning is required (*54*). Discriminant analysis (*55*) is a common multivariate statistical approach for data classification (for example, in 2016, 3,107 scientific articles were published on the use or improvement of discriminant analysis – search terms used in Web of Science: discriminant analysis, in title/keywords/abstract).

The simplest forms of DA are linear (LDA) and quadratic (QDA). Linear Discriminant Analysis can be seen as a regression line whose orientation divides a high-dimensional space, reducing the dimensionality while keeping each class separate from the other classes. In practice, the optimal orientation is the one that minimizes the within-class variance and maximizes the between-class variance (*56*). The main assumption of LDA is that all the classes have a common variance-covariance matrix, i.e. the relationships between classes and explanatory variables are independent from class membership, while the differences between classes are dependent only on the mean.

When the variance-covariance matrices are not homogeneous for two or more classes, linear discriminant analysis cannot be applied. Instead the QDA can be employed. The QDA discriminant function is:

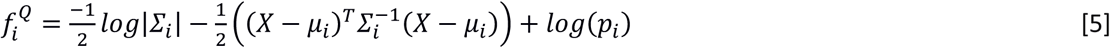

where **X** is the matrix of variables, *μ* the vector containing the mean of each variable and **Σ** is the variance-covariance matrix, and p_i_ the “prior” probability of each point to belong to the class *i. i* is the subscript for class *i*, with *i* = 1, …, N where N is the total number of classes.

*f* is calculated based on a training dataset (class memberships are known). The larger the *f* value, the higher the probability that the point belongs to that group. For a training dataset, the *p_i_* can be calculated in several ways, usually by “equal priors” method: each class has a prior probability equal to 1/N. In this analysis, and in order to take into account the spatial proximity of the classes, a local frequency prior method was used. It estimates the class *p_i_* prior probability as the relative frequency of *i* labels in the neighbourhood. Similarly, predicting a label for a new point means looking at the local proportion of each class (as classified from the training dataset) around the new point.

Once *f* is maximised with the training dataset, new data points can be classified by calculating *f* for the new point and for each class (equivalent to calculating the position of a point with respect to all available class centroids), and assigning to it the class index at which corresponds the maximum *f*.

#### QDA algorithm for optimising the number of classes and classification

The QDA has been embedded into an algorithm that determines the optimal number of ecological classes and their geographic delimitation for each area.

The algorithm steps are the follows:

i. Define the initial number of classes, N_0_. The initial choice has been N_0_ = the number of land cover classes in the area. This decision is made on the assumption that mosquito distribution is significantly predicted by land use and land cover. The co-variates are all the environmental variables described above. *Splitting algorithm*
ii. QDA is applied to N_0_ classes in the first iteration, otherwise to N_j_.
iii. The class with lowest probability is then split into two sub-classes of similar size based on the criterion of minimum intra-class variance. At the iteration, *j*, the number of classes is N_j_ = N_j-1_+1.
iv. Repeat ii and iii until N_j_ is equal to a maximum number of classes, here fixed to subjectively to 8. *Merging algorithm*
v. Set *j*=1
vi. Starting from N_0_, merge the two classes with the largest probability that members belong to both classes. At the iteration, *j*, the number of classes is N_j_ = N_j-1_-1.
vii. Apply QDA to N_j_ classes
viii. Repeat vi and vii until N_j_ is equal to a minimum number of classes, here fixed to 2. *Selection of the optimal number of classes*
ix. The optimal number, N*, of classes is selected based on the Wilk’s criterion (*57*). The largest reduction in the Wilk’s criterion between two consecutive classes (equivalent to a sharp decline below the trend in the graph plotting Wilk’s Lambda on the y-axes and number of classes in the x-classes) indicates the optimal number of classes. *Classification*
x. In the final step, all the points are classified in one of the N* classes. Uncertainty is measured as the probability that a point belongs to any of the other classes.

The Wilks’ criterion is based on the following general equation:

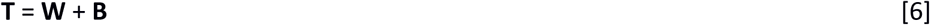

where **T** is the total sums of squares and products matrix, **W** is the total sums of squares and products within groups and **B** is the total sums of squares and products between groups. The Wilks’ criterion or Wilks’ Lambda (*L*) is the ratio of the determinants of **W** and **T**:

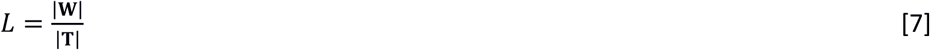

therefore, minimizing *L* is equivalent to minimising |**W**|.

The analysis was carried taking all the environmental variables at their original spatial resolution, and providing the output (classification) at 30m resolution (the same as the land cover resolution).

### Spatial allocation of the sample households

Locations of the sampling points, in each sampling site (Figure 1), follow a “lattice plus close-pairs” design (*58*) which combines regular lattice (efficient for predictions) and random points as close pairs (efficient for parameter estimation) (*59*).

For an easier understanding of the sampling design, we refer to the six locations distributed in West and East Africa as sampling sites (Fig. 1). Each sampling site will contain *M* sampling points. Each sampling point contains *V* households. Therefore the total number of households sampled per sampling site is *M* × *V*.

For all sites except Migori, the lattice plus close pairs design is realised under two conditions: (i) 70% of sampling points are in lattice and 30% are distributed randomly (as usually applied in simulation analyses, i.e. (*58*) and (*60*)) (Figure 2); (ii) each stratum must contain a number of points proportional to the stratum size (*61*):

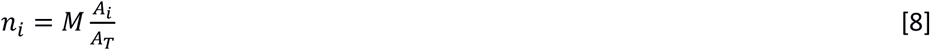

where *n_i_* is the number of points for class *i*; *A_i_* the area of class *i*; and *A_T_* is the total area. The term “close pairs” here is used loosely, since not all the points in the grid will have a close pair, and some close pairs may be shared between points.

**Figure 2.**
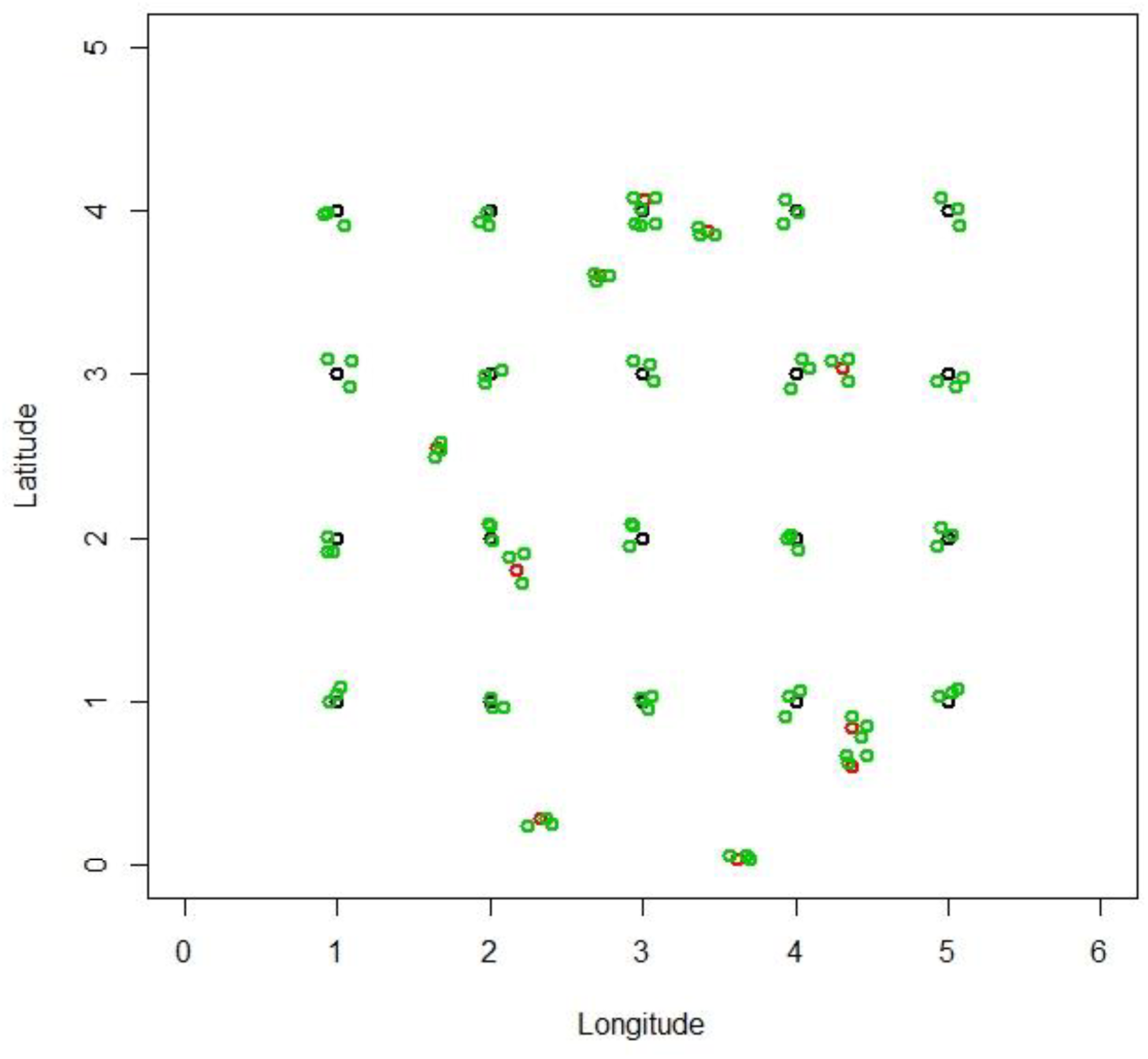
Example of lattice with close pairs design adopted in this work. Black dots, sampling locations in regular grid; red dots, sampling locations allocated randomly; and green dots are the households identified sufficiently close to the sampling locations (V). Plot was made using R-cran 3.5.0 (http://r-project.org).

In Migori alone (where a previous sampling campaign, AIRS, took place) an adaptive sampling design was trialled in which AIRS sampling served to inform the location of the *M* sampling points. From the LGCP model (see above), we estimated the prediction variances at each grid cell, and attributed the *M* locations to the cells with highest prediction variance (*27, 62-64*).

### Effect of stratification on sample size and improvement of mosquito abundance models

In order to evaluate the effect of stratification:

a. on the ratio between the Poisson rate parameter of mosquito counts from a survey (λ_1_) and the Poisson rate parameter of mosquito count from a sub-sample of it (λ_2_);
b. and on the goodness of fitting of mosquito abundance models;

we have considered a mosquito sampling campaign from Uganda. This data is from 104 health subdistricts (HSD) where estimates of *An. gambiae* and *An. funestus* densities (as determined by a standard collection method) for both male and female mosquitos are available. The sampling design was based on a CRT with ten houses selected at random from each HSD, and mosquito collection was made every six months for two years (http://www.isrctn.com/ISRCTN17516395).

For objective (a) we have employed a Poisson exact text on the null hypothesis that the ratio between λ_1_ (obtained from the entire Uganda mosquito collection data) and λ_2_ (obtained from a sub-sample of mosquito collections of the Uganda data) is equal to 1, i.e. the two conditional means are not different (*65*). The test is performed by first randomly sampling 2, 3, 4 and 5 mosquito collection locations from each strata. For each of these sample sizes the λ_2_ is calculated and the test performed. The process has been repeated 999 times, to randomise the location selection, and 95% confidence interval from the all bootstrapping are estimated. The procedure above was then compared with a sampling design that randomly extracts the same amount of locations from the entire dataset but independently from the strata to which they belong.

For objective (b) we fitted the total number of mosquitoes for each species and at each location (over the two years of collection) using the ecological strata produced by performing the same methodology described in the above section “Stratification”. The model fitting employs a Poisson generalized linear model (*65*) and model comparison against the null model is performed using a MANOVA test (*18*).

## Results

### Sample size

We estimated the parameters associated with the spatial autocorrelation of the AIRS mosquito surveillance. The maximum likelihood estimation of the Log-Gaussian Cox Process parameters returned: intercept of 21.77, spatial variance (sill – i.e. the amount of variance dependent on distance) of 14,478, spatial range of 16 km (i.e. the maximum distance at which variance increases with distance) and 0 nugget (variance independent of distance that can be due to measurement errors or un-explained factors). Therefore, according to the model, all the variation is considered to be spatially-dependent up to a 16km range. The Matern kappa parameter (shape parameter) was 1.5 (Supplementary Information 1).

In order to estimate the impact of the sample size on model fitting and predictions, we simulated a Log Gaussian Cox Process with known covariance function (the Matern in Supplementary Information 1). The results are reported in Table 1.

**Table 1.** Total variance in the parameters of the Gaussian process (intercept, sill, nugget, range) and standard errors for the predictions at different sample sizes.

| Sample size | Total variance in the parameter of the Gaussian process | Standard error in predictions |
| --- | --- | --- |
| 15 | 8034 | 194 |
| 30 | 4726 | 93 |
| 75 | 454 | 44 |
| 150 | 71 | 31 |
| 200 | 14 | 22 |
| 300 | 0.86 | 9 |

With 30 sampling locations, the prediction error and the total variance in the LGCP parameters was halved compared to 15 locations, and 20 times less when using 75 locations. The standard error in predictions is the maximum number of mosquitoes predicted in excess or in deficiency to the true mean. Therefore with 30 traps it is estimated a maximum error of 93 mosquitoes around the real mean and with 200 locations an error of 22 mosquitoes.

In order to improve local estimates, more than one household for each sampling point can be employed. Therefore if we take two households for each of the 30 sampling points, the total number of households is 60. The effect of the use of more than one household in model fitting and prediction is shown in Table 2.

**Table 2.** Total variance in the parameters of the Gaussian process (intercept, sill, nugget, range) and standard errors for the predictions at different number of households at each sampling point, with 30 sampling points.

| Number of households at each sampling point | Total variance in the parameter of the Gaussian process | Standard error in predictions |
| --- | --- | --- |
| 2 | 5207 | 71 |
| 3 | 2073 | 61 |
| 4 | 1851 | 22 |
| 5 | 701 | 19 |
| 6 | 605 | 19 |
| 7 | 118 | 18 |

With four households and 30 sampling points we expect the same prediction error as using 200 random sampling points distributed across the entire area and each containing a single household (comparison of standard errors in Tables 1 and 2) but higher variance in the parameters. Using between five and seven households has little impact on the standard error in the predictions, although there is a significant improvement in the model fitting as the number of households increases (see total variance column in Table 2). Consequently, the sampling design was chosen with 30 locations and four households which was considered a good balance in terms of standard errors, model fitting and economic feasibility.

### Ecological classification

The ecological classification identified two classes for Migori, Obuasi, Muleba and Aboude; three classes in Malindi and four in Grand Popo. The Wilk’s criterion, measured as Wilk’s Lambda, for Malindi is shown in Figure 3; for other sites, they are provided in supplementary information files. The biggest improvement (i.e. largest decrease of the Wilk’s Lambda) is in the change from 2 to 3 classes (Figure 3). For the other sites, see Supplementary Information 2.

**Figure 3.**
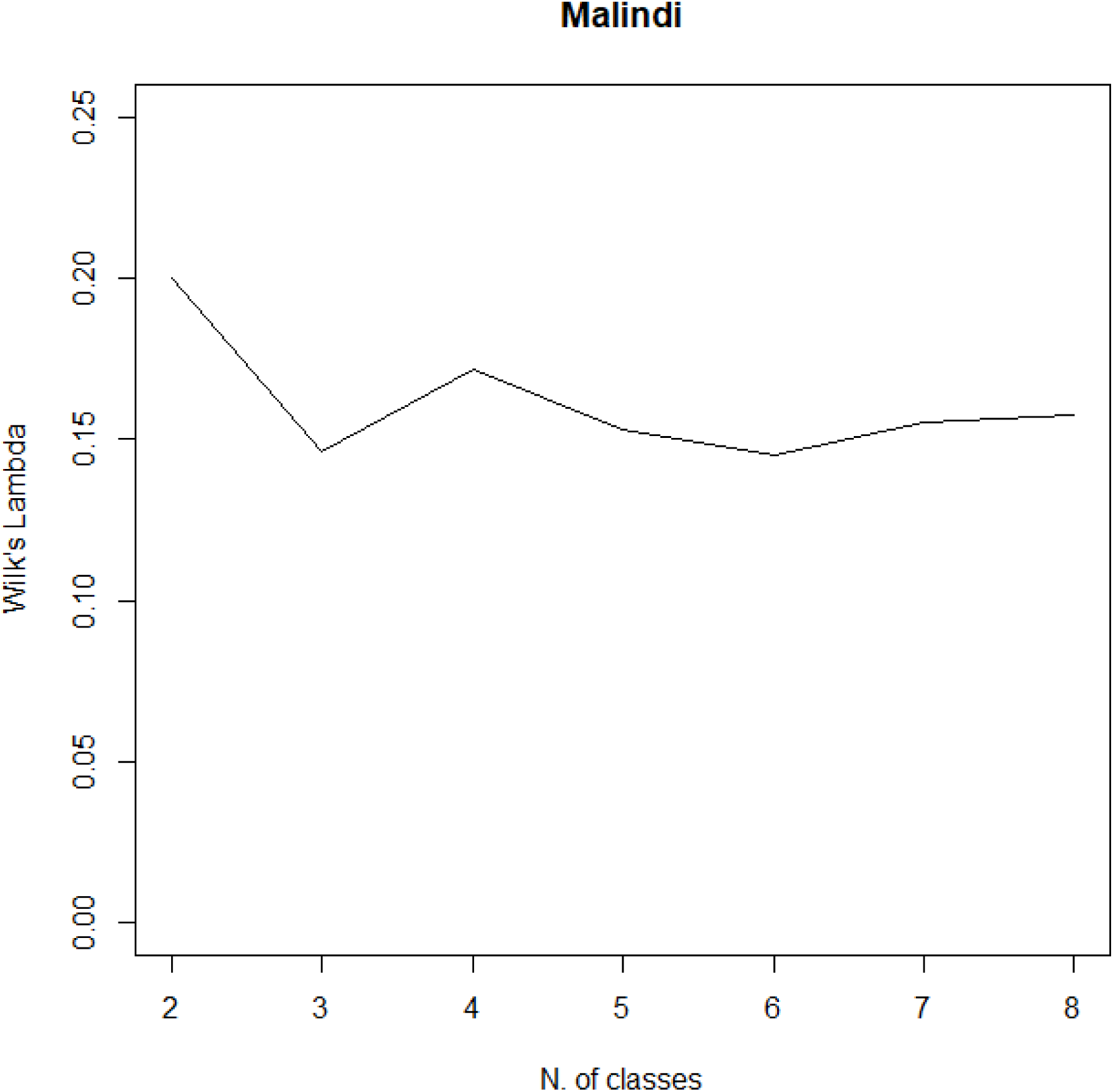
Wilk’s Lambda criterion for Malindi. Graph was made using R-cran 3.5.0 (http://r-project.org).

A hierarchical numerical classification of the sites and classes is shown in Figure 4. This dendrogram was obtained from the agglomerative method (classes are aggregated into progressively larger groups) group average (*66*). The latter accounts for the average distances or similarities between all the members of the new group and those of the others.

**Figure 4.**
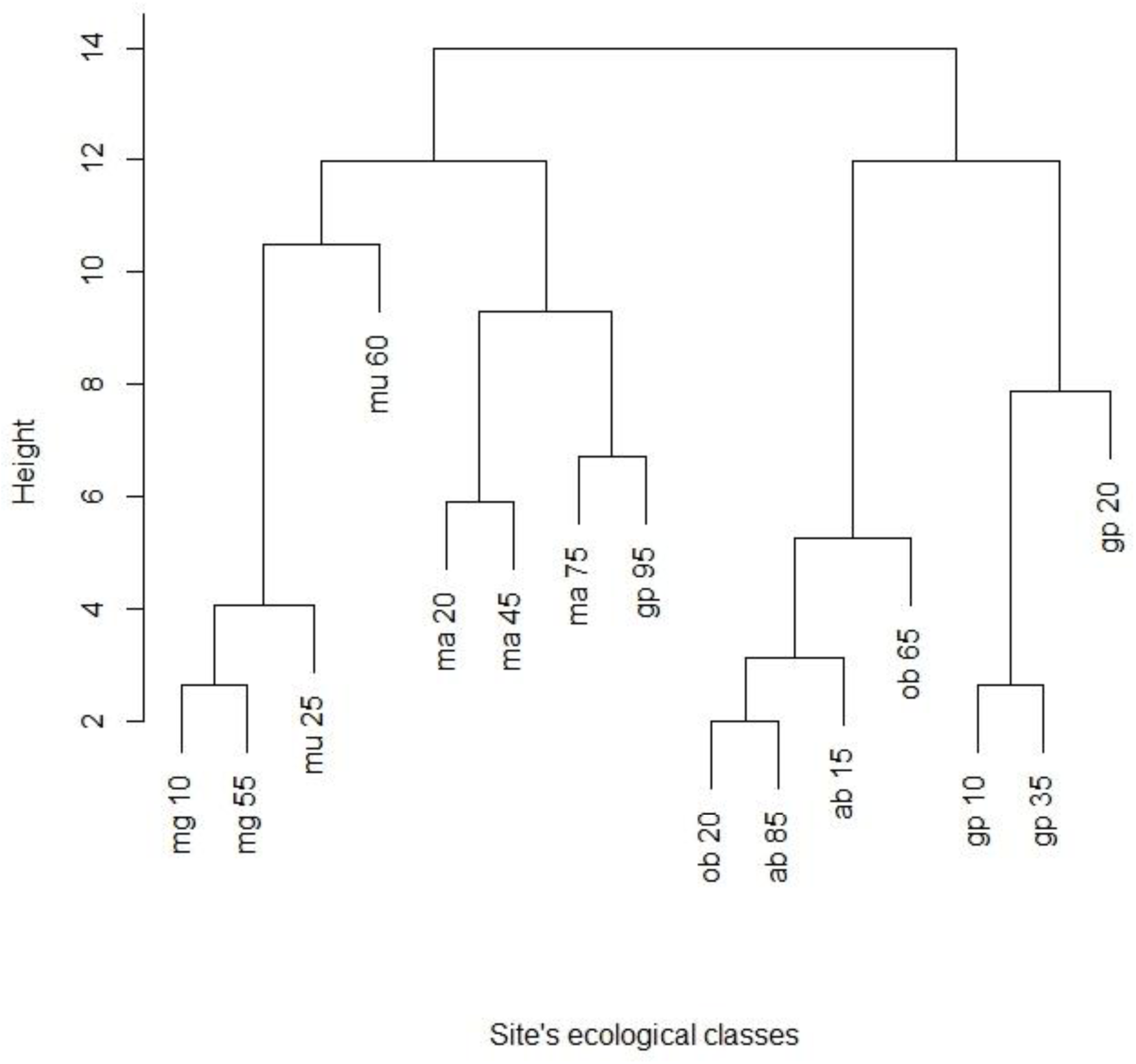
Dendrogram of agglomerative hierarchical clustering of the ecological zones. mg, Migori; mu, Muleba; ma, Malindi; gp, Grand Popo; ob, Obuasi; and ab, Aboude. For the class number see Table 3. Graph was made using R-cran 3.5.0 (http://r-project.org).

**Table 3.** Classes delineated by ecological classification: description, location and colour used in the maps.

| Class | Description | Colour |
| --- | --- | --- |
| 10 | **Cultivated land. Medium** ET mean and variance, EVI mean, variance and amplitude, Temp mean, variance and amplitude, Precipitation; **High** ET amplitude and Elevation. | Mango |
| 15 | Mixture of **Cultivated land** and **Forest. Low** ET mean, EVI variance and Temp mean. **Medium** EVI mean, elevation and Precipitation. | Red |
| 20 | **Forest. Medium** ET mean amplitude and variance, EVI mean, variance and amplitude, Temp mean and variance, Elevation and Precipitation; **High** Temp amplitude. | Dark Green |
| 25 | Mixture of **Cultivated land, Shrubland** and **Wetland. Low** ET mean, EVI variance and Temp mean. **Medium** Elevation and Precipitation. | Orange |
| 35 | Mixture of **Shrubland** and **Grassland. Low** Temp amplitude and Precipitation; **Medium** ET mean and variance, EVI mean, variance and amplitude, Temp mean and variance; **High** ET amplitude. | Light green |
| 45 | Mixture of **Shrubland, Grassland** and **Wetland. Low** EVI amplitude, Temp variance and Precipitation; **Medium** ET mean and amplitude, EVI mean and variance, Temp mean and variance; **High** Temp mean and ET variance. | Pink |
| 55 | Mixture of **Urban, Forest, Wetland** and **Grassland. Low** Temp mean; **Medium** ET mean, EVI mean and variance, and Precipitation; **High** Elevation. | Yellow |
| 60 | **Wetland. Low** ET mean, EVI mean and variance; **Medium** Temp mean and Elevation; **High** Precipitation. | Navy |
| 65 | Mixture of **Urban, Tundra, Wetland, Water bodies** and **Grassland. Low** ET variance and amplitude; **Medium** Elevation; **High** ET mean, EVI mean, variance and amplitude, Temp amplitude and Precipitation. | Purple |
| 75 | Mixture of **Water bodies** and **Urban. Low** ET mean, EVI mean and amplitude, Temp variance and Precipitation; **Medium** ET amplitude and EVI variance; **High** ET variance and Temp mean. | Sky |
| 85 | Mixture of **Grassland, Water bodies** and **Urban. Low** ET variance and amplitude, and Elevation; **Medium** Temp mean; **High** ET mean, EVI mean, variance and amplitude, Temp variance and Precipitation. | Brown |
| 95 | Mixture of **Wetland, Water bodies** and **Urban. Low** EVI mean and amplitude, and Temp amplitude; **Medium** ET mean, EVI variance, Temp variance and Precipitation; **High** ET variance and amplitude, and Temp mean. | Grey |

\*Low, values lower than 25% quartile; Medium, values between 25% and 75% quartiles; High, values larger than 75% quartile.

The heights in Figure 4 represent the dissimilarities between classes, which are very small for some intra-location comparisons (Migori, Kenya, mg10 and mg55; Grand Popo, Benin, gp10 and gp35) and indeed some inter-country comparisons (ob20 in Obuasi (Ghana) with ab85 in Aboude (Cote d’Ivoire)). The classification suggests geographic homogeneity for most of the sites, since locations aggregates more than classes. In the four groups ((i) mg10/mg55/mu25/mu60, (ii) ma20/ma45/ma75/gp95, (iii) ob20/ab85/ab15/ob65, and (iv) gp10/gp35/gp20), Migori in Kenya and Muleba in Tanzania (321km apart) are clustered together, as are Obuasi in Ghana and Aboude in Cote d’Ivoire (300km apart); while Malindi (with exception of class 95 in Grand Popo), whose closest location is Migori at 666km, and Grand Popo, which is closest to Obuasi at 384km, form their own clusters. Both Grand Popo and Malindi are coastal sampling locations, albeit on opposite sides of the African continent.

Figure 5 shows the ecological classification and its uncertainty for the Malindi area. The same maps for the rest of the sites are shown in Supplementary Information 3.

**Figure 5.**
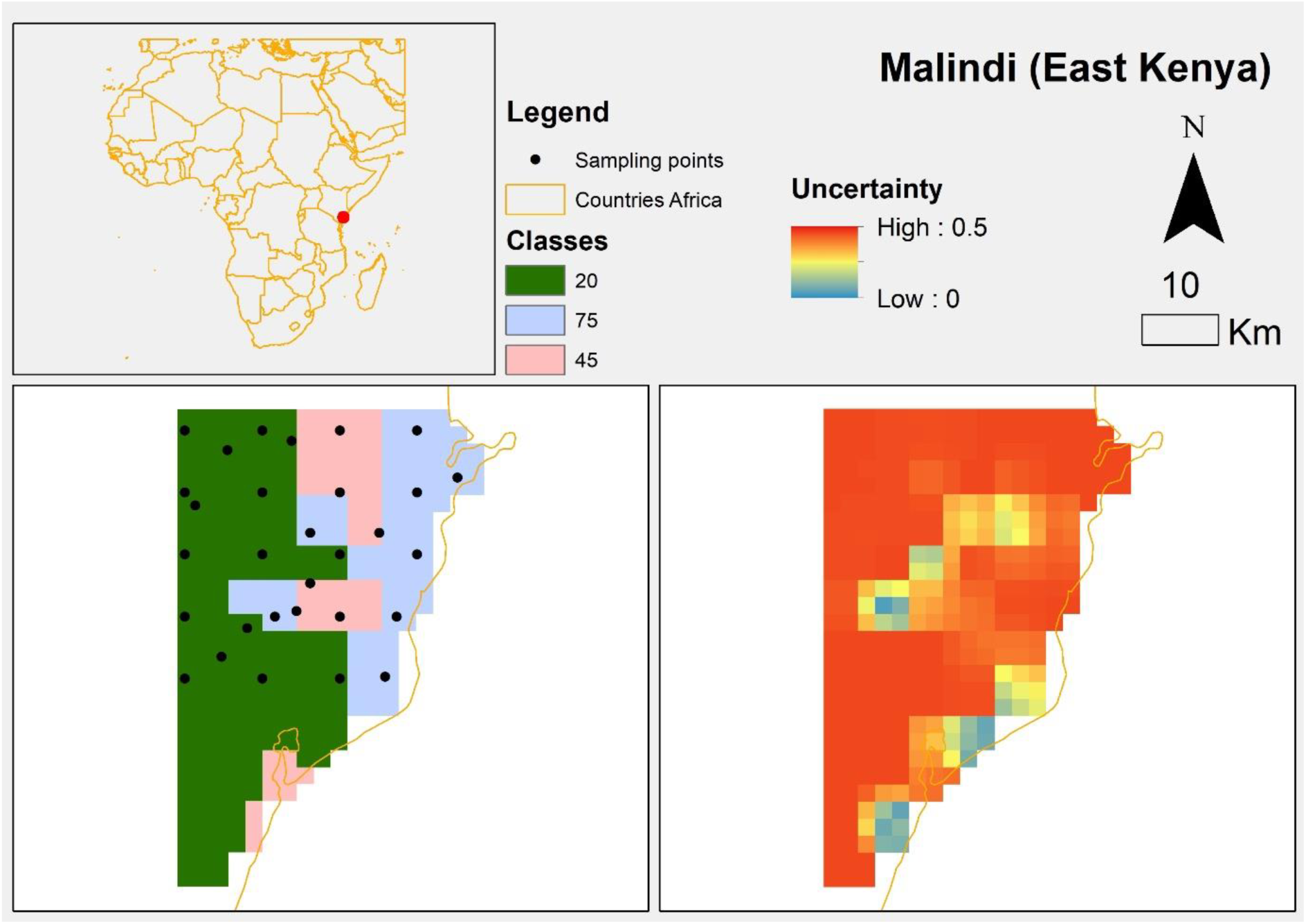
Ecological classification and uncertainty for the area of Malindi. Map was made using ArcMap 10.4 (http://desktop.arcgis.com/en/arcmap/). Source administrative limits: http://www.maplibrary.org/library/index.htm.

### Sample locations

In Figure 5 the sampling locations for Malindi are shown overlaid on the ecological classes. These locations are obtained from the lattice with close pairs sampling design, with a batch of 4 households (not shown in the figure) at each sampling point. In this lattice with close pairs sampling design 20 of the 30 locations are deployed in a 4 × 5 regular grid, and the rest allocated randomly as described in the methods. This general objective was modified to weight the number of locations by ecological classes; thus some of the points in the grid may have been adjusted slightly to be contained in the new class, though never by more than half of the grid-cell size.

In all the sites, each of these 30 locations constitutes a cluster of 4 households as in Figure 2.

The sampling location in Migori (Supplementary Information 3) followed an adaptive sampling design, in which only class 10 was sampled because of the constraint that previous samples (AIRS) only targeted this class. The allocation of 30 samples and households in class 10 in Migori were based on the prediction variance, i.e. new sampling points are allocated at the centre of 30 pixels with largest prediction variance (*64*) (Figure 6).

**Figure 6.**
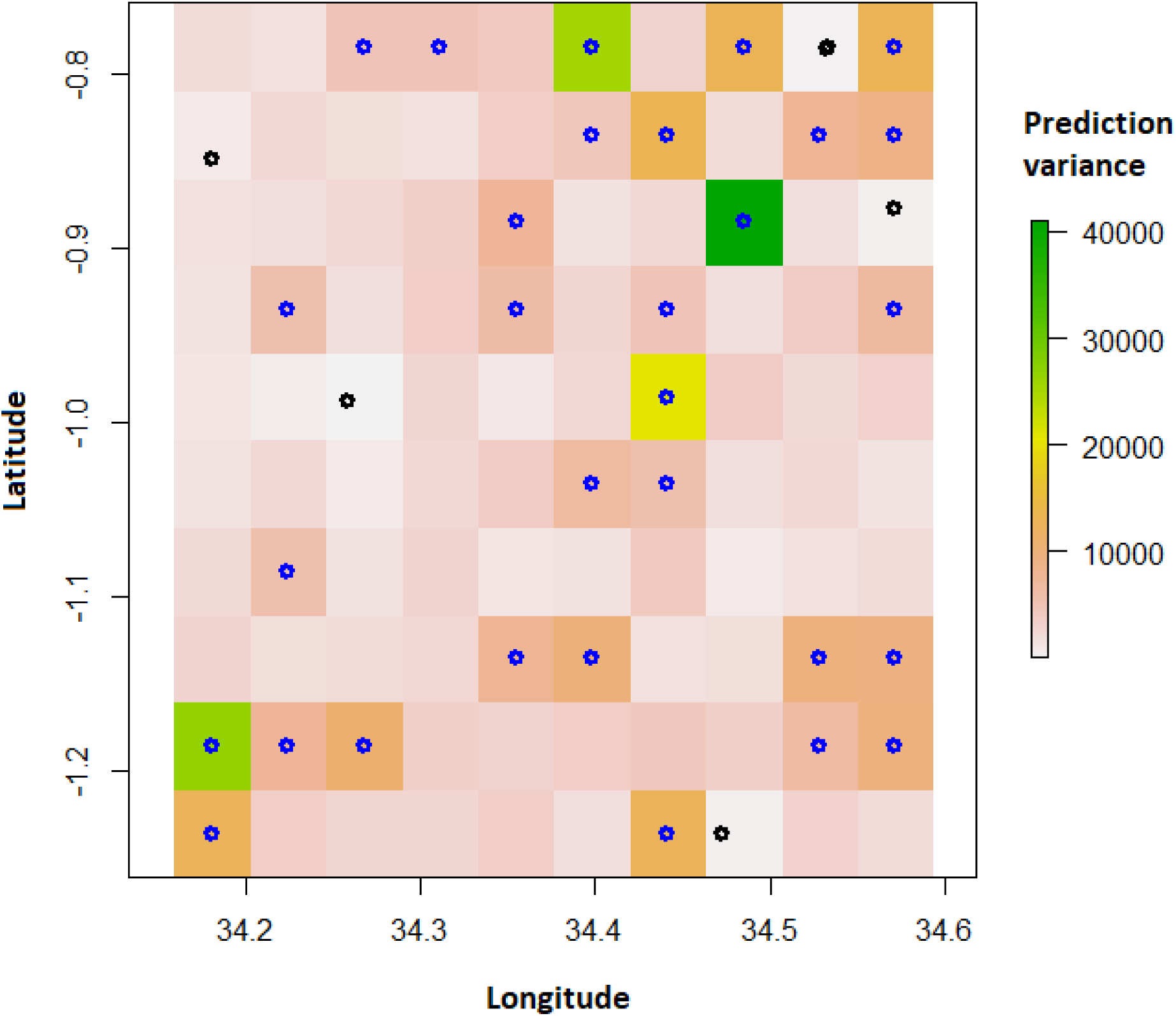
Adaptive sampling for Migori in class 10. Black dots are the AIRS mosquito surveillance locations. The Blue dots are the adaptive locations, which are targeting the cells with largest prediction variance (represented with green, yellow or orange colours). Graph was made using R-cran 3.5.0 (http://r-project.org).

### Effect of stratification on sample size and improvement of mosquito abundance models

For this objective we have used the Uganda dataset (see methods), which contains a large mosquito sampling campaign (104 districts, 1040 households) carried out over for 2 years. The first step was to identify the ecological strata for each cluster (of households) location. By applying the same stratification method described for the GAARDian project (see methods) we identified 4 ecological zones.

The stratification was first used to evaluate if a sub-sample (up to 20% of the full dataset) of the mosquito collections, extracted from each strata, is still representative of the full mosquito collection obtained in Uganda.

From Table 4 confidence intervals show that the stratification produces ratios that are not significantly different from 1 (i.e. λ_1_ and λ_2_ are not significantly different) for any sample size. In contrast for random sampling all ratios between the two rate parameters are significantly higher than 1 for all the sample sizes. Finally Table 5 shows that stratification improves the model fitting of mosquito counts when compared to a null model.

**Table 4.** 95% Confidence interval (CI) of the rate ratio between λ_1_ and λ_2_ (Poisson distribution rate parameters from mosquito counts in the full survey and mosquito counts in a sub-sample of locations respectively), where λ_2_ is calculated for each sample size. The sample size refers to each strata for a total of 2*4, 3*4, 4*4 and 5*4 locations, where 4 is the number of strata. In the case of complete random sampling (last 3 rows), then the 2*4, 3*4, 4*4 and 5*4 are the number of locations sampled independently from the strata.

|  | CI | 2 | 3 | 4 | 5 |
| --- | --- | --- | --- | --- | --- |
| Stratified | 0.025 | 0.94 | 0.91 | 0.86 | 0.85 |
|  | 0.5 | 1.05 | 0.99 | 0.93 | 0.9 |
|  | 0.975 | 1.16 | 1.08 | 1 | 1.07 |
| Random | 0.025 | 1.11 | 1.11 | 1.03 | 1.04 |
|  | 0.5 | 1.25 | 1.21 | 1.11 | 1.07 |
|  | 0.975 | 1.4 | 1.33 | 1.21 | 1.12 |

**Table 5.** ANOVA analyses of Poisson generalised linear models for female (F), male (M) and total (F+M) mosquitoes of *An gambiae* and *An. funestus*. Residual deviance is in % of the Null deviance.

| Species | Sex | Residual deviance | P value |
| --- | --- | --- | --- |
| *An. gambiae* | F | 87 | 2.2 e-16 |
|  | M | 89 | 2.2 e-16 |
|  | F+M | 88 | 2.2 e-16 |
| *An. funestus* | F | 88 | 2.2 e-16 |
|  | M | 93 | 2.2 e-16 |
|  | F+M | 89 | 2.2 e-16 |

## Discussion

Vector-borne disease control and monitoring rely on vector surveillance, mostly carried out using trap-based indices and, more recently, remote sensing data (*13, 24*). Trap-based indices (density, population changes, distribution, etc.) are calculated from mosquito catches and require a system of traps dispersed in the field in sufficient numbers to represent mosquito population ecology and dynamics. Conversely, remote sensing data can be used to define the ecological level of disease risk based on mosquito ecological suitability (*61, 67*). This is a cheaper and quicker option, but may not have the spatial and temporal resolution necessary for practical interventions (*23, 68-71*). In a sampling design, trap-based indices and remote sensing analyses must be seen as complementary tools, since remote sensing data contain the information necessary to define location and distance between sampling points.

Sample size, location, estimator and strategy (i.e. model based or not, adaptive or not) are the fundamental characteristics of a study design (*25*), which affect the likely success of describing the studied process (e.g. disease or organism distribution and abundance), its stochasticity, and, consequently, the accuracy of the estimates.

Ecologists are now equipped with algorithms, open information (*72*) and datasets that enable a better understanding of the biology and spatial distribution of populations, which allows optimization of collection site placement to best describe natural processes. Ecological/environmental classification is now possible for every region in the world (*68*). Failure to exploit these data in ecological and genomic sampling frameworks ignores the spatial variability of favourable, unfavourable or neutral habitats, therefore random or transect sampling designs may or may not be representative of the ground conditions and characteristics (*73*). Even a grid design can be biased towards larger ecological classes and may miss linear features (i.e. a river passing between collection points) (*1*). The consequence of which could be an over- or under-estimate of the true abundance, even when the population phenology is correctly delineated (*73*).

In the illustrative example presented we demonstrate how these approaches may be used to develop an *a priori* sampling strategy to sample malaria vectors for genomic and ecological studies. The ecological classification presented for each site returned a maximum uncertainty ranging from 0.37 to 0.44 depending on the site (Figure 5 for Malindi and Supplementary Information 3 for the other sites), which can be interpreted as the probability that a grid node belongs to a different class. This level of uncertainty shows that classification identified dominant classes, in other words each point was allocated to an ecological class with an absolute minimum probability of at least 0.56. In addition, the ecological classification also shows that areas (the 6 sampling sites) with putatively the same land cover are still ecologically different when considering the full set of environmental variables (temperature, precipitation, elevation, evapo-transpiration and vegetation), and that geographical proximity is a dominant factor in ecological clustering (see also (*74, 75*)) (Figure 4). It is therefore not surprising that sites cluster much more strongly within country than within ecotype; e.g. forest (class 20, Figure 4) in Malindi (Kenya) is not equivalent to forest in Obuasi (Ghana) or Grand Popo (Benin). Ecological classification, while not often used in modelling mosquito populations and communities for medium- and large-scale analyses, represents a complex interaction of the environmental and socio-economic conditions (*23*).

Another factor that we accounted for during our sampling design is the spatial autocorrelation of mosquito catches (model-based sampling design) (*12*). The effect of strong autocorrelation can reduce the overall statistical power (and the overall biological significance of the study) as it results in effectively a lower sample size (because the assumption of independence is violated), underestimates of variance, and increases in type I error (*10*). Geostatistical approaches, such as the one applied here (*8, 62*), can lead to unbiased estimates of population parameters and avoid the risks and limitations of random, or haphazard, selection of sampling locations (*10*). Given the requirements to satisfy both parameterization and predictions (*59*), the simulated inhibitory design adapted from (*62*) in order to contain clusters of households at each sampling point, has shown that with 120 sampling houses for each site distributed across 30 sampling points, we achieve the same prediction error (main goal) as from 200 points allocated at random, albeit at the expense of parameter accuracy. However, there were important limitations in the sample size/location calculation. Firstly, they are based on limited pre-existing mosquito surveillance data from Migori, which may not describe the different spatial scales of the mosquito abundance distribution (*24*). This is a concern due to the large variation in abundance levels observed throughout the period, but that can be solved by deploying an adaptive sampling design, i.e. concentrating the new samples where we have the largest uncertainties (our knowledge is poor) in the process of interest (abundance or a level of abundance). The advantages are an improvement of the estimates with a lower number of samples (*76*). In addition, we are assuming that the mosquito population dynamics in Migori are similar to those in the other sites. The ecological classification has the advantage of correcting for local mosquito population dynamics although this is not a full solution. Ideally, Migori could have been used to analyse the effect of the ecological classification on mosquito estimates. Unfortunately, the pre-existing surveillance samples are located in the same ecological zone (Supplementary Information 3) making it impossible to simulate the effect of the ecological classification on the sample size/location optimization (Diggle *et al*. 2010). For this reason we evaluated the effect of stratification on sample size using a different dataset (Uganda). This shows that 10 to 20% of mosquito collections randomly selected from strata are representative of the full survey. This result shows that stratification can be applied at any stage of the sampling campaign, and even if it was not considered at the initial (planning) phase, it can adaptatively inform the subsequent sampling phases or collections and optimise the sampling costs (subsequent sub-sampling of each strata).

In addition, using the Uganda dataset, we have also shown that the ecological stratification improves model fitting, again representing a model feature that can be applied in both pre-analysis and post-analysis of sampling campaigns.

An element not considered in this analysis but that requires discussion is the temporal frequency and length of the sampling campaign. Designs for temporal sampling raise the same challenges as spatial designs, along with additional considerations. These include: “*is it better to trap six times in each of two houses, or twice in each of six houses, or four times in each of three houses? And in the latter case, is it necessary that the nights should be at weekly intervals, or would the easier task of sampling over four consecutive nights yield a similar amount of information? Should the same ‘fixed’ houses be sampled on each occasion, or should a new set be chosen randomly on each occasion?*” (extracted from (*77*)). Answering these questions requires relatively lengthy longitudinal studies and a knowledge of *Anopheles* population dynamics. Fortnightly collections are common in mosquito sampling designs (*78*), and enable cost-effective descriptions of seasonality and variation in mosquito abundance (*18*). On the other hand, positioning traps during peaks of mosquito abundance can significantly overestimate the rate of population increase and the level of abundance (*73*), and only sampling over two or more years may accurately account for cyclical fluctuations in vector abundance (*77*).

Our analysis provides an example of how to fully describe the assumptions, conditions and constraints of sampling strategies. We do not expect other researchers to precisely replicate our methodology, e.g. the use of 4 houses in 30 sampling locations depends on previous abundance analysis but this may change when more information will be available (adaptive sampling), but we have shown how open-data sources and ecological information can be implemented in the initial steps of sampling design. Our literature review shows that the specifics of sampling design are poorly reported and we therefore suggest that even when sampling is based on expert-opinion decisions, a full description of the sampling design should be provided to make the sampling repeatable or comparable or usable for subsequent similar studies. For example, field constraints such as presence of the disease or vector or host, vegetation type and density, elevation, field hostility, logistic feasibility, potential interference, human proximity, breeding sites, and risk of trapping material theft (*13, 20, 21, 24*), which often are the major influence in the sampling design need to be declared and described. In fact, previous sampling campaigns are often used to inform future sampling design, and therefore standardization of sampling designs and protocols are now a priority (*12*).

In conclusion, big and open data and research outputs could enhance the power of ecological and genomic studies (*3*), facilitating the growth of complex and multidimensional algorithms. In the specific field of vector biology and genomics, there is an urgent need to establish standards for mosquito sampling design and description in scientific reports. One of the first steps is to facilitate training and workshops (*11*) but also the improvement of publishing standards (i.e. requiring authors to fully disclose the sampling design) in order to produce a collection of high quality and usable sampling designs along with their results.

## Acknowledgments

We thank Prof Peter Diggle and Dr Michael Chipeta for the useful advices on the presented analyses. This study was supported by funding from the Medical Research Council, Grant Number MR/P02520X/1

## Authorship

LS, EL, DW and MD conceived the study. LD, AE, AE-Y, BK, JM and EO provided background information and data on the sampling sites and inspected the areas. LS performed the statistical analysis. LS, EL, DW and MD drafted the manuscript and all the authors contributed to its final version.

## Data accessibility statement

All data used in this research is available through the links described in the paper. Any additional data or R codes can be requested from the corresponding authors (LS).

